# Parallel Wires: A Conserved Principle of Contralateral-Ipsilateral Segregation in the Visual Corpus Callosum

**DOI:** 10.1101/2025.09.30.679451

**Authors:** Jiaowen Wang, Yanming Wang, Yiping An, Shishuo Chen, Benedictor Alexander Nguchu, Huan Wang, Muhammad Mohsin Pathan, Yueyi Yu, Sinan Yang, Ying-Qiu Zheng, Yang Ji, Hao Wang, Yifeng Zhou, Bensheng Qiu, Xiaoxiao Wang

**Author notes:** These authors contributed equally to the work (Jiaowen Wang, Yanming Wang, Yiping An, Shishuo Chen).

## Abstract

How the brain maintains distinct information streams within dense white matter is a fundamental question. We investigated whether the visual corpus callosum transmits information via segregated “parallel wires” or mixed pathways. This distinction is critical: a mixed architecture would render the signal’s origin ambiguous, whereas a segregated arrangement implies that spatial position tracks the direction of information flow. Using high-field fMRI and Bayesian modeling in humans, we demonstrate a segregated architecture featuring distinct contralateral and ipsilateral channels. This functional segregation mirrors a precise anatomical arrangement in mice, where dual-color viral tracing and light-sheet microscopy reveal that callosal axons remain spatially segregated in distinct laminae after crossing the midline. Our findings establish a conserved “parallel wires” principle of callosal organization, providing a new framework to decode directional information flow and assess pathway-specific damage in neurological disease.

## 1. Introduction

The corpus callosum (CC), a structure highly conserved within mammalian brains ^1,2^, integrates information between the two hemispheres of the brain ^3,4^. Within the visual system, the CC tracts fuse inputs from the contralateral visual cortices ^5^ into a single, coherent perception ^6,7^. Our study focuses on the forceps major (FMA), the largest component of the visual callosum ^8^, where a question remains: is visual information transmitted via segregated parallel streams or mixed?

Adjudicating between these models is critical because it addresses a fundamental limitation of functional Magnetic Resonance Imaging (fMRI). An fMRI signal detected within a white matter bundle is directionally ambiguous, as its hemisphere of origin is unknown. A ‘mixed’ architecture, where axons from both hemispheres intermingle, would make this origin impossible to resolve. Conversely, a segregated ‘parallel wires’ architecture—where tracts from each hemisphere occupy distinct spatial territories—would mean that the signal’s spatial location directly reveals its origin. Confirming a segregated parallel architecture would therefore represent a fundamental leap, moving beyond mapping static links (’who is connected’) ^9–11^ to decoding dynamic information flow (’who is talking to whom’). This principle is analogous to laminar fMRI, which leverages spatial segregation across cortical layers to differentiate processing streams ^12–14^.

A fundamental principle of neural wiring is that axons maintain the topographic information of their source regions: visual callosal connections, for instance, exhibit a visuotopic organization mirroring their cortical origins ^15,16^. Historically, testing this principle within white matter has been challenging. However, recent advances have demonstrated that reliable fMRI signals can be detected in white matter ^17–21^ and visuotopic organization is preserved within these tracts ^22,23^. This allows the “segregated versus mixed” question to be reframed as a directly testable hypothesis: a ‘segregated’ architecture predicts voxels with singular visuotopic representation, whereas a ‘mixed’ architecture predicts voxels with multiple visuotopic representations.

To directly test this hypothesis, we employ a multi-modal, cross-species investigation. In humans, we combine ultra-high-field 7T fMRI ^24^ with Bayesian population receptive field (pRF) modeling ^25^ to map the functional topology, and use diffusion tractography ^26,27^ to delineate its anatomical substrate. For mesoscale validation, we employ dual-color neuronal viral tracing and whole brain light-sheet microscopic imaging ^28,29^ in mice to reveal the underlying circuit architecture. This multi-modal approach allows us to construct the comprehensive functional blueprint of the primary interhemispheric visual pathway.

## 2. Results

### Bayesian pRF Modeling Reveals Distinct Receptive Field Topologies in FMA

To directly test the ‘parallel wires’ versus ‘mixed pathway’ hypotheses, we characterized the visual field representations within the forceps major (FMA). We utilized the minimally preprocessed 7T retinotopy fMRI dataset from 178 participants of the Human Connectome Project (HCP) ^24^. The functional data was non-linearly normalized to the standard template, and averaged to create a single group-averaged fMRI dataset for subsequent analysis. We applied the Bayesian pRF modeling ^25^ to frame the question into a model comparison: a segregated architecture would manifest as voxels best explained by a single pRF, whereas an integrated architecture would prefer models with multiple pRFs. The Bayesian model comparison decisively favored a segregated architecture. The single pRF model, which posits that callosal voxels carry information from one visual hemifield, was overwhelmingly dominant (expected probability = 0.92). Conversely, models assuming integration (dual or triple pRFs) received minimal support (Fig. 1A).

**Figure 1.**
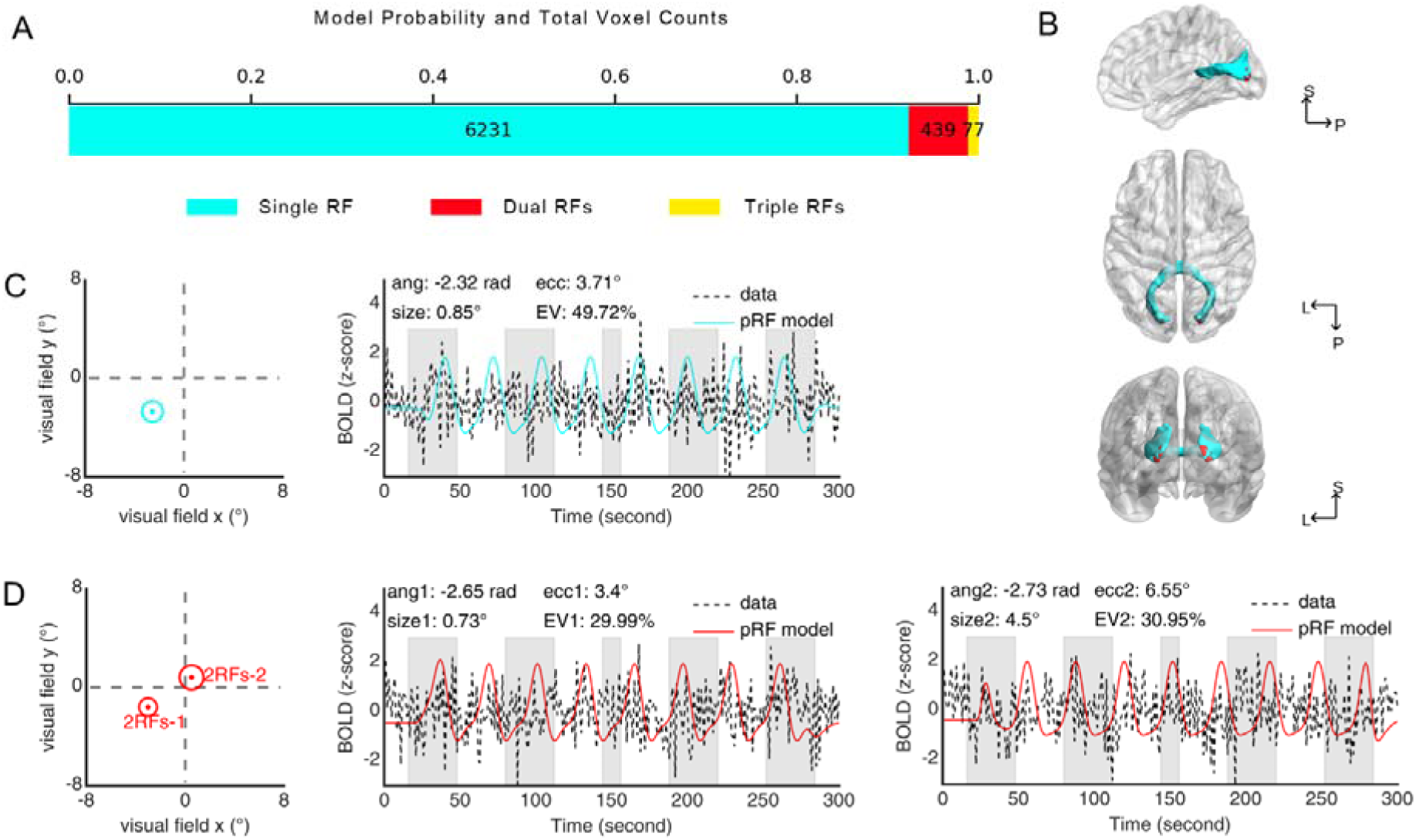
Mapping of The Visual Fields on FMA Voxels Based on Single, Dual, and Triple Receptive Field Models. **(A)** The mapping of the visual fields was performed based on single, dual, and triple pRF models, with expected probabilities or voxel counts of 6231 (blue) for single, 439 (red) for dual, and 77 (yellow) for triple pRF models. **(B)** Spatial distribution patterns show that voxels best explained by the single pRF model (blue) populate the core of the FMA fiber bundle, whereas the sparse population of integrative dual pRF voxels (red) clusters near the occipital gray-white matter interface. **(C, D)** Representative voxels for the single (blue) and dual (red) models. The left panels indicate the pRF center (dot) and size (circle) in degrees of visual angle (dva). The right panels display the time series fitting, where the model predictions (red solid lines) robustly capture the observed BOLD signal variance (black dashed lines), confirming the high fidelity of the functional data. EV: explained variance.

The spatial distribution of these models was highly structured. Voxels best explained by the single pRF model were distributed throughout the core of the FMA fiber bundle. In contrast, a small cluster of voxels, particularly those near the interface of the gray and white matter at the occipital cortex, aligned with dual and triple pRFs models, (Fig. 1B). The pRF models robustly captured the Blood Oxygenation Level Dependent (BOLD) signal variance (Fig. 1C-D), confirming the high fidelity of our functional data.

### Parallel Contralateral–Ipsilateral Organization of the FMA

Polar angle mapping reveals that the FMA is dominated by contralateral representations, which account for 68.65% of voxels. The remaining 31.35% form a distinct ipsilateral stream running parallel to the main tract. This parallel contralateral-ipsilateral organization has not been reported previously, and therefore, this study establishes a new paradigm for understanding how visual information is represented or processed by the corpus callosum or the FMA. Our newly discovered parallel architecture in FMA received strong support from analysis of the functional gradient, where parallel spatial segregation, similar to that observed in pRF mapping (in polar angle maps), was evident (Fig. 2B). Further analysis to confirm the dominance of the contralateral organization in the parallel architecture was performed, where the gradient cortical projection indeed demonstrated the presence and dominance of contralateral representations on the visual cortex (Supplementary Fig. S1).

**Figure 2.**
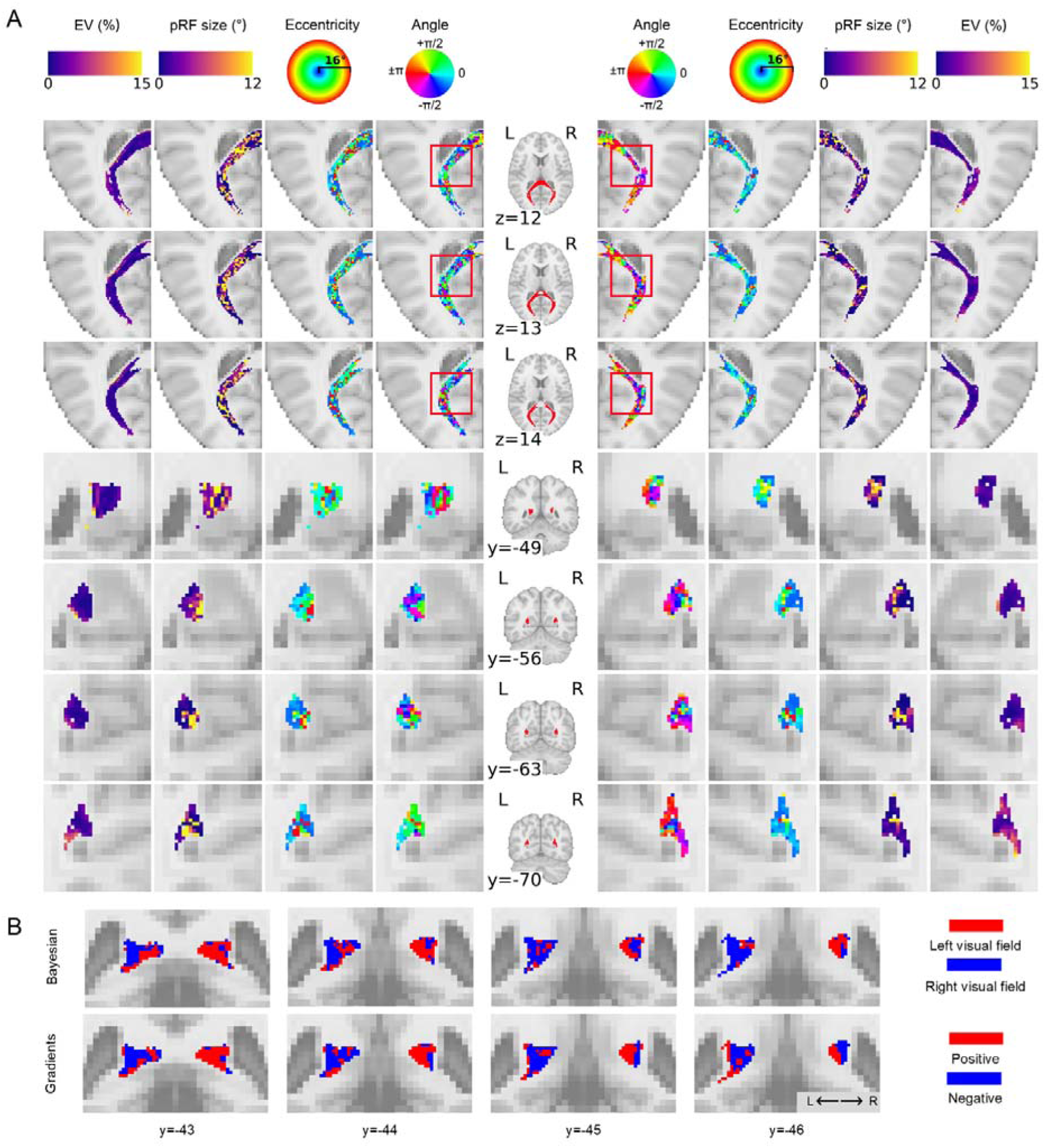
Parallel Contralateral–Ipsilateral Organization of the FMA. **(A)** Population receptive field (pRF) parameter maps for the winning single pRF model, shown in selected axial (top rows) and coronal (bottom rows) views. From the center outwards, columns display the FMA mask, polar angle, eccentricity, pRF size, and explained variance (EV). Polar angle topographies demonstrate a striking parallel organization: dominant contralateral representations (red/orange for left visual field, cyan/blue for right) are spatially segregated from, yet run alongside, distinct ipsilateral streams (highlighted in red box). Eccentricity and pRF size maps further characterize these streams, showing a preferential representation of the visual periphery with large receptive fields. **(B)** Cross-methodological validation confirms this angular organization in a representative coronal slice. Functional gradient analysis (bottom) delineates the same parallel retinotopic structure observed in the Bayesian pRF estimates (top), with consistent segregation of left (red) and right (blue) visual field representations. All maps are overlaid on the ICBM 152 2009c standard brain template.

Further characterization of these streams revealed a clear retinotopic profile. Eccentricity maps (Fig. 2A, eccentricity columns) showed a preferential representation for the foveal and para-foveal visual fields (Supplementary Fig. S2B). As expected, pRF size (Fig. 2A, size columns) scaled linearly with eccentricity (R²=0.999, p<0.001; Supplementary Fig. S2C). The hemodynamic responses were delayed and prolonged compared to gray matter (Supplementary Fig. S2D, peak latency: 8.81 s vs. 5.00 s), consistent with white matter physiology ^22,30^. The robustness of these findings was affirmed by split-half reliability tests, which revealed high consistency of pRF parameter estimates (polar angle ICC=0.4538; Supplementary Fig. S3).

To ensure these findings were not merely an artifact from group-averaging, we repeated our analysis on four randomly selected subjects. Although the results from the four subjects were noisier than those derived from the whole dataset, they still confirmed the presence of contralateral-ipsilateral organization (Supplementary Fig. S4, particularly in the coronal view), with contralateral dominance. Their analysis of tractography on callosal pathways at the dorsal level also validated these findings, where a similar organization was observed, characterized by a dominance of single, contralateral-preferring receptive fields alongside distinct ipsilateral representations (Supplementary Fig. S5). These results were consistent with laminar organization, and together they demonstrate high reproducibility for our findings even at the individual-subject level.

### Contralateral Dominance Persists Within Dual pRF Voxels

For voxels responding to a dual pRF model, structured patterns or representations were observed on both sides of the brain (Supplementary Fig. S6A). While these voxels responded to stimuli in both hemifields, they exhibited a strong contralateral bias, contralateral pRFs explaining significantly more variance than the ipsilateral pRFs (Supplementary Fig. S6B). Furthermore, even when dual pRFs shared a contralateral alignment, their receptive fields remained spatially separate (Supplementary Fig. S6C, D). However, the entire integration process follows a contralateral-ipsilateral organization, with contralateral processing of visual information dominating the information flow in the overall architecture of visual field processing.

### Fiber Tractography Delineates the Anatomical Substrate of Functional Streams

To delineate the anatomical substrates of the functionally-defined visual field representations within the FMA, we employed a deterministic fiber tracking approach at the group level ^26^. The fiber tractography of the FMA, performed using seeds derived from locations that responded to single and dual pRF models (Fig. 3A-B), unraveled two parallel fiber bundles traversing the FMA. The two fiber bundles were spatially and anatomically distinct, characterized by a clear topological separation throughout their tracts (Fig. 3C-D), supporting a model of functional segregation mediated by anatomically discrete pathways.

**Figure 3.**
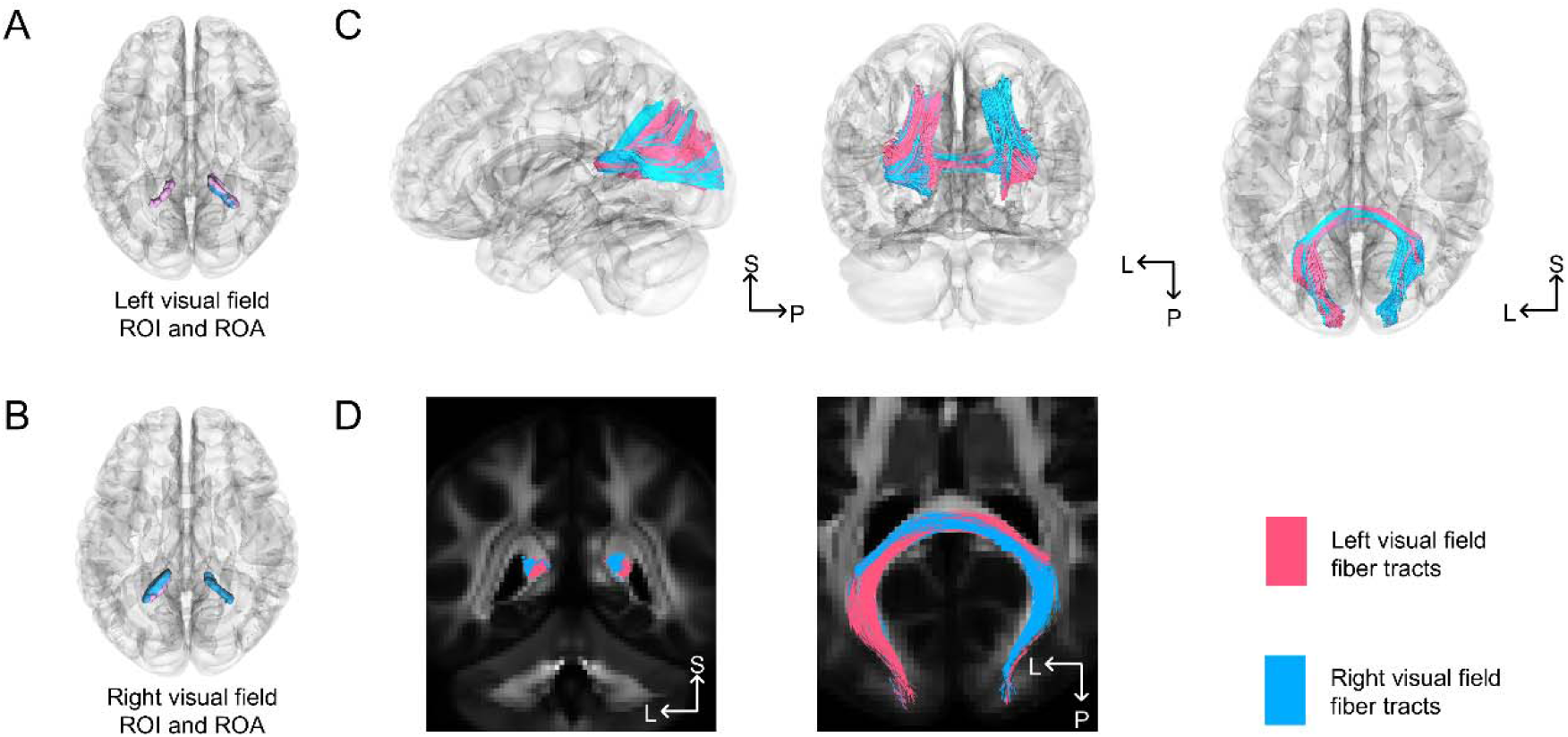
Deterministic fiber tracking demonstrating parallel information flow in the FMA. **(A, B)** Tractography seeding strategy based on functional pRF maps. To isolate specific information channels, regions of interest (ROI, pink) and regions of avoidance (ROA, blue) were defined for the left **(A)** and right **(B)** visual field representations, respectively. **(C)** Whole-brain reconstructions in MNI space reveal that these functionally distinct streams correspond to two anatomically separate fiber bundles. **(D)** Cross-sectional coronal and axial views further demonstrate this topological organization, where fibers carrying left visual field information (pink) and right visual field information (blue) maintain a strict parallel, non-overlapping trajectory throughout the fiber bundle.

### Anatomical and Mesostructural Evidence for Segregated Pathways

To investigate the mesoscopic architecture underlying these functional streams, we employed dual-color viral tracing and high-resolution light-sheet microscopy in the mouse brain (Fig. 4A). Selective labeling of pathways from the left and right primary visual cortex (V1) confirmed robust contralateral projections (Fig. 4B-C), establishing a clear anatomical basis for the contralateral dominance observed in the human fMRI data. Crucially, this dual-color tracing revealed a precise spatial organization that mirrors the functional topology observed in humans. At the commissural midline, axonal populations from the two hemispheres exhibited a transient convergence, forming a narrow zone of partial overlap (visualized as yellow in Fig. 4E). This localized interdigitation offers a compelling structural basis for the sparse population of integrative dual-pRF voxels identified at the gray-white matter boundary. Distal to this midline crossing, however, the two fiber populations diverged and resumed a parallel trajectory, resolving into spatially distinct bundles within the contralateral hemisphere (Fig. 4D, E). This robust post-commissural segregation, which was quantified by significant spatial separation (Dice = 0.26±0.10, Jaccard = 0.15±0.07, mean±st.d.), provides the anatomical substrate for the overwhelmingly dominant population of single-pRF voxels. This segregation of anatomical structures can be seen clearly through magnification (see Fig. 4F-G, Supplementary Fig. S7A-G). The two anatomical structures form a laminar-like arrangement, with axon projections from one hemisphere localized dorsal to the other (Fig. 4F-G, Supplementary Fig. S7A-G). Collectively, these mesostructural findings establish a principle of dominant anatomical segregation punctuated by localized midline integration, thereby providing a definitive validation for the functional architecture identified with human fMRI.

**Figure 4.**
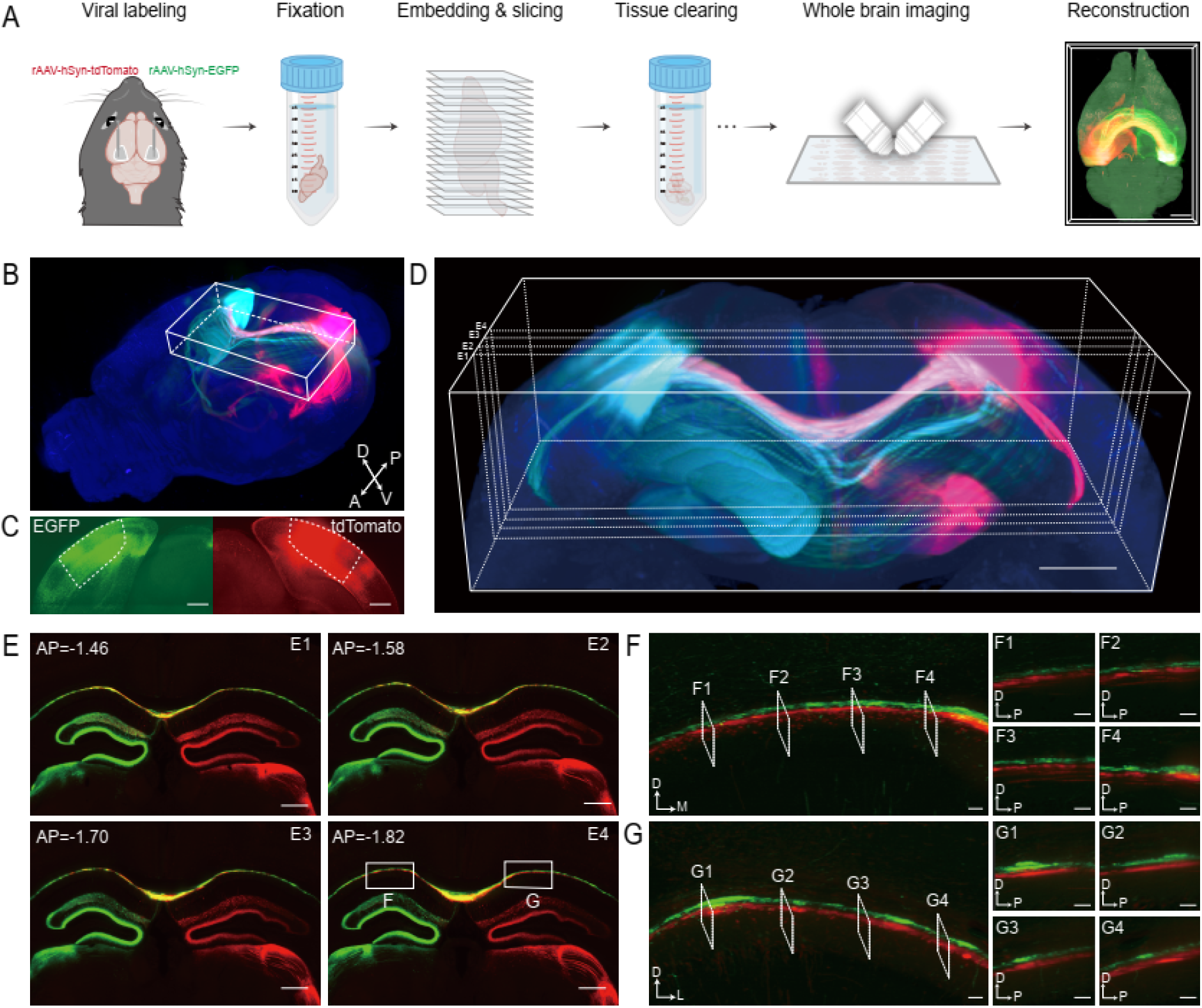
Laminar segregation of contralateral axons in the mouse visual corpus callosum. This figure presents mesoscale anatomical evidence from the mouse brain that validates the principle of segregated functional pathways observed in human fMRI. **(A)** Pipeline of whole brain viral labeling and imaging: viral labeling, brain fixation with transcardial perfusion, brain embedding and slicing, tissue clearing, whole-brain imaging and 3D reconstruction. Scale bar: 1 mm. **(B)** Three-dimensional reconstruction and visualization of the axonal projections expressing rAAV-hSyn-EGFP (green) and rAAV-hSyn-tdTomato (red) in the whole brain. A: anterior; P: posterior; D: dorsal; V: ventral. **(C)** Confined viral expression in the respective V1 injection sites confirms the specific labeling of these two distinct hemispheric pathways. Scale bar: 300 μm. **(D)** The ROI showing axonal projections in the CC, with a volume of 3950 × 1440 × 1970 μm^3^. Scale bar: 500 μm. **(E)** Coronal sections reveal a dynamic organizational principle. At the midline of the CC, the axons from both hemispheres intermingle, producing a transient zone of overlap (yellow). As the bundles project laterally away from the midline, they resolve into clearly separated red and green populations. Scale bar: 500 μm. AP: anterior-posterior. **(F, G)** High-magnification views of the regions outlined in (E1-E4) reveal the precise internal organization of the callosal bundles. The main panels show coronal sections, and the corresponding sagittal sections (F1-F4, G1-G4) taken along the medial-lateral axis demonstrate a definitive laminar segregation. Axons from one hemisphere (green) are consistently positioned dorsal to axons from the other (red). Scale bars: 50 μm.

## 3. Discussion

This study establishes that the visual CC is governed by a conserved principle of contralateral-ipsilateral segregation. Our multi-modal evidence reveals that this tract is not a passive mixer, but a structured pathway where information flow is strictly organized into parallel streams. In humans, functional mapping demonstrated that callosal channels are dominated by inputs from a single hemifield, a finding corroborated by both pRF modeling and functional gradient analysis (Fig. 2B) and aligned with discrete anatomical tracts (Fig. 3). We then confirmed the conserved nature of this organization at the mesoanatomical level in mice, where dual-color viral tracing revealed a novel, non-overlapping laminar arrangement of axons from the contralateral and ipsilateral hemispheres (Fig. 4). These convergent findings reveal that the visual CC operates as a set of parallel, segregated conduits, where each pathway independently processes information from one side of the visual field, thereby preserving the spatial origin of the signal within the white matter.

### From Human Function to Mesoscale Anatomy

A longstanding challenge in neuroscience is bridging the macro- and meso-scales of brain connectivity, especially when the focus of the study involves integrating human non-invasive imaging with animal invasive tracing ^31^. At the macroscale, diffusion MRI tractography is the primary tool for mapping the human connectome ^32^. At the mesoscopic scale, researchers have established viral tracing, considered the anatomical gold standard ^33^, to provide the ground truth. This has been dramatically enhanced by technological innovations in high-resolution, whole-brain imaging ^28,29,34^ and the creation of comprehensive connectivity atlases ^35^. The established approach, therefore, is to use this definitive meso-anatomical data to validate and constrain the large-scale pathways inferred from human tractography ^36–38^. However, this structural validation is inherently limited, as diffusion MRI is blind to the direction and nature of information flow.

Our study operates within this validation paradigm but extends it into the functional domain. Building on foundational work that validated white matter BOLD signals ^17,19,39,40^, we use pRF modeling to move beyond signal detection and decode the informational content of callosal pathways. This allows us to reverse the conventional workflow, i.e., we employ viral tracing and light-sheet microscopy not to confirm a structural pathway, but to uncover the anatomical substrate behind the functional principle that has been discovered in our first in vivo functional analysis. The cross-species approach implemented here is deeply grounded in the conserved developmental and molecular blueprint of the corpus callosum ^2,41,42^, which shows similarity in both mice and humans, despite differences in their scales and cognitive demand ^43^. While our findings establish segregation as a conserved principle, the human visual CC is likely more complex than the mouse’s. Therefore, future investigations in non-human primates, which bridge this evolutionary gap, will be essential for a complete understanding of the visual callosal pathways.

### Integrating White Matter with Laminar-Resolved Function

Bridging the gap between microscale circuit models and human cognition remains a central goal in neuroscience. By leveraging ultra-high-field imaging to resolve activity across cortical depths, laminar fMRI has emerged as a powerful tool in this endeavor. This technique allows for the in-vivo testing of canonical circuit models, for instance by dissociating feedforward, input-driven activity in middle cortical layers from feedback signals in superficial and deep layers ^12–14^. Our discovery of information segregation within white matter provides a critical new dimension to this framework. The combination of anatomically segregated white matter pathways with functionally specific cortical laminae presents a powerful synergy for human brain circuit research. A question emerges, therefore, whether this segregation represents a general organizing principle that extends to other commissural and intrahemispheric tracts.

This capacity to resolve distinct information channels is of immediate relevance to clinical neuroscience. A significant innovation of our approach is its ability to delineate directional structural connectivity. For instance, following a unilateral stroke or traumatic brain injury, it becomes possible to assess not just if the CC is damaged, but precisely which directional pathways are compromised—either the pathways projecting from the lesioned hemisphere, or those projecting to the other, or both ^44,45^. This anatomical specificity is also poised to refine emerging functional measures like GM-WM connectivity analysis ^46–48^. By providing the structural blueprint of parallel conduits, our work enables the decomposition of a single tract’s functional signal into pathway-specific components, a critical step for developing more sensitive biomarkers for conditions ranging from psychiatric disorders ^20,49^ to individual cognitive traits ^50^. Although this study leveraged the high signal-to-noise ratio of 7T MRI, the rapid maturation of high-resolution fMRI sequences on 3T platforms promises to democratize this approach, accelerating its translation from basic science to widespread clinical application ^51^.

### Developmental Origins of Segregated Callosal Architecture

The bundled architecture of the corpus callosum is established during development by a precise interplay of molecular and mechanical forces ^52^. During this period, cell adhesion molecules (CAMs) mediate the homophilic “zippering” of axons into cohesive fascicles ^53^. Several repulsive cues confine the bundling processes. These cues include the interaction between Ephrin-A4 on callosal axons and EphA receptors in the surrounding tissue, which establishes a “permissive corridor” that prevents axonal migration and steers the entire tract across the midline ^1,54^. However, while these established mechanisms of fasciculation and guidance account for the formation of the callosal projection as a whole ^42,55^, they do not fully explain the highly organized internal structure we report. The subsequent refinement processes that establish the precise laminar segregation of ipsilateral and contralateral axons remain a key area for future investigation.

### The Observation of Dual pRFs

The observation of dual pRFs is an expected consequence of signal pooling ^56^, where a single fMRI voxel averages signals from thousands of axons. In the FMA, this reflects the known convergence of bilateral visual inputs near the vertical meridian, a phenomenon well-documented at the cortical boundaries ^57–59^. We therefore interpret these dual pRFs as the macroscale signature of localized zones where contralateral and ipsilateral fibers intermingle. Crucially, the sparse and localized nature of these integrative voxels, in contrast to the widespread distribution of single pRFs, reinforces our primary conclusion: the visual callosum operates predominantly as a set of parallel, segregated information conduits.

## 4. Summary

In summary, this study establishes a conserved “parallel wires” principle of callosal organization, resolving a fundamental question regarding the transmission of information across the brain’s largest white matter tract. By integrating ultra-high-field human fMRI with mesoscale viral tracing in mice, we demonstrate that the visual corpus callosum avoids signal mixing in favor of a strictly segregated architecture. In humans, this manifests as parallel functional streams where dominant contralateral inputs remain spatially distinct from ipsilateral channels; in mice, this is mirrored by a precise, non-overlapping laminar segregation of axons. These findings fundamentally reframe our understanding of interhemispheric communication, moving beyond static connectivity to reveal a dynamic, directionally specific highway where spatial location encodes the origin of information. This framework provides a novel blueprint for decoding directional information flow in the living human brain, offering critical precision for future basic and clinical neuroscience research.

## 5. Methods

### 5.1. Subjects

The subjects included 178 young adults from the HCP (109 females, 69 males), aged 22 to 35. All participants underwent 7T retinal topology fMRI and DTI data. All subjects had normal or corrected-to-normal visual acuity. Each subject was assigned a six-digit HCP ID.

### 5.2. HCP Datasets

Complete retinotopy datasets (six fMRI runs) were acquired for a total of 181 subjects (109 females, 72 males), aged 22–35, as part of the Young Adult HCP (https://www.humanconnectome.org/study/hcp-young-adult/datareleases). To ensure data completeness and quality, we excluded 3 individuals with incomplete imaging records required in this study. The final cohort consisted of 178 participants (109 females, 69 males) with fully available 7 T retinotopic functional MRI (fMRI) and diffusion MRI (dMRI) data, which were used for subsequent analyses. The 7T retinotopy dataset includes four packages: (1) Retinotopy Task fMRI Functional Preprocessed Extended; (2) Structural Preprocessed for 7T, with the same spatial resolution and space as the fMRI data; (3) Diffusion Preprocessed, with 1.05 mm spatial resolution diffusion and structural data in native space. Stimuli for the retinotopy dataset comprised colorful object textures windowed through slowly moving apertures. Apertures and textures were generated at a resolution of 768 × 768 pixels and were constrained to a circular region with a diameter of 16.0 dva. The apertures included clockwise or counterclockwise rotating wedges (RETCW, RETCCW), expanding or contracting rings (RETEXP, RETCON), and moving bars (RETBAR1, RETBAR2). For each stimulus type, a 5-minute fMRI scan was conducted, consisting of 300 volumes. The stimuli were presented in the following order: RETCCW, RETCW, RETEXP, RETCON, RETBAR1, and RETBAR2.

### 5.3. Preprocessing

We utilized the 7T retinotopy dataset from the Human Connectome Project (HCP), for which the data collection and standard preprocessing pipelines are extensively described elsewhere ^24,60,61^. The minimally preprocessed data provided for each subject included volumetric fMRI time series in MNI space (1.6 mm isotropic, TR=1s), high-resolution T1-weighted (T1w) structural images (0.7-mm isotropic), and diffusion-preprocessed data, including fractional anisotropy (FA) images.

To improve the accuracy of spatial normalization, particularly for T1w’s limited contrast within the white matter, we implemented a custom registration procedure that leverages both T1w and FA images. This procedure began by individually intensity-normalizing the T1w and FA images for each subject to their respective global means. Subsequently, a composite structural image (T1w+FA) was generated for each subject by summing these two normalized volumes; this process was repeated for the MNI template’s T1w and FA images to create a corresponding template composite. Nonlinear registration parameters were then computed by registering the individual’s T1w+FA composite to the template composite using the Advanced Normalization Tools (ANTs) package ^62^. In the final step, these derived transformation parameters were applied to the subject’s original 1.6 mm fMRI data to bring them into the 1.0 mm MNI template space with high anatomical fidelity.

Prior to group averaging, several processing steps were applied to the time series of each individual subject. For each retinotopy run, the initial 8 volumes were discarded to ensure T1 signal stabilization. The remaining time series of each run was then normalized (e.g., to percent signal change) and linearly detrended to correct for baseline shifts and signal drift. The cleaned runs for each subject were then concatenated in their original acquisition order. Finally, a single group-averaged fMRI dataset was created by averaging these fully preprocessed time series across all subjects for subsequent analyses.

### 5.4. Regions of Interest (ROIs)

A mask for the FMA was derived from the standard library provided by XTRACT, a component of the FMRIB Software Library (FSL). XTRACT offers a comprehensive set of predefined masks for white matter tractography, facilitating consistent and reproducible tract-based analysis ^63^. The library and additional details can be accessed at https://fsl.fmrib.ox.ac.uk/fsl/fslwiki/XTRACT. The parietal CC connection mask was derived from Radwan et al. ^64^. The whole-brain WM mask was generated using the automated segmentation tools provided by the FMRIB Software Library (FSL) ^65^. To avoid signal contamination from neighboring GM voxels, the FMA white matter mask was expanded inwards by 4 voxels. It is worth noting that the probabilistic maps were in MNI152 space. In this study, the T1 structural image in MNI152 space was registered with the asymmetric T1 image from the ICBM152 2009c template to generate a conversion file. Subsequently, the FMA fiber probabilistic tracking template from XTRACT was registered to the functional image space to extract the corresponding fMRI time series.

### 5.5. A Combined Model of pRF and Hemodynamic Response Functions (HRFs)

The Bayesian pRF framework allows the specification, estimation, and comparison of receptive field models, seeking the optimal balance between model accuracy and complexity. As Zeidman et al. described ^25^, neural activity at time t is modeled using a multivariate normal probability density function N. The response was summed over each illuminated pixel on the screen and scaled by a parameter. Structurally like the classical pRF approach, the Bayesian pRF framework comprises two components: a neuronal model describing the cortical response to visual stimulation, and a hemodynamic model accounting for neurovascular coupling and the resulting BOLD signal. The neural parameters and the elements of the matrix are limited to a specific range of values through latent variables. These variables are utilized to compute the neuronal response. The specific parameter settings and model specification are comprehensively described below. The Bayesian pRF model employs a series of spatial coordinates *p_i_* to represent the set of stimulus positions *U_t_* at time *t*.

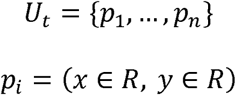

The difference-of-Gaussians (DoG) model, which incorporates center excitation and surround suppression, is expressed as the difference between two Gaussian profiles. Here, the neural response z(*t*) is modeled as the sum of responses to all stimulus positions under the difference between two an Gaussian profiles, each weighted by the multivariate normal probability density function *N*(*p_i_*|μ, Σ): excitatory center with gain *β_c_*, center μ, and covariance Σ*_c_*, and an inhibitory surround sharing the same center but with gain *β_s_* and covariance Σ*_s_*. Here, μ denotes the two-dimensional pRF center and Σ the covariance matrix determining receptive field size.

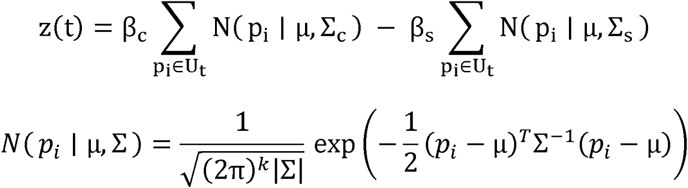

Latent variables *I_p_* and *I_θ_* were introduced to parameterize the pRF center in polar coordinates (ρ,θ) By applying the normal cumulative distribution function (NCDF), the center position (μ*_x_*, μ_y)_ was rigorously constrained to remain within the circular stimulus region across all optimization steps.

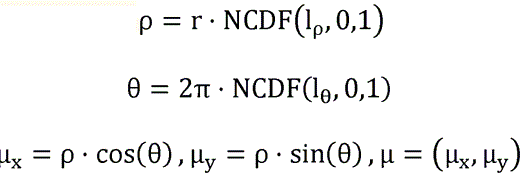

The size of the pRF was parameterized by its standard deviationσ, with the range constrained by the latent variable *l_θ_*, and subsequently transformed within the model. Here, *r* and *r_0_* denote the radius of the circular stimulus region and the minimum allowable pRF size, respectively. The excitatory covariance matrix Σ*_c_*, and inhibitory surround Σ*_s_* yields the DOG circular model.

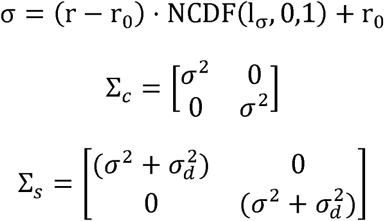

β*_d_* and σ*_d_* ensure positive center activation and negative surround response, enabling the model to capture both excitatory and suppressive components of receptive field organization.

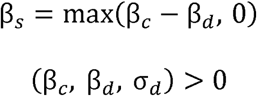

Zeidman et al. fed the predicted neural activity described above into the extended Balloon model to obtain the predicted BOLD response. In this study, the time series prediction was obtained by convolving *z*(*t*) with a model of the hemodynamic response function (HRF; *h*(*t*))^66,67^, which is composed of two gamma functions.

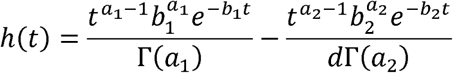

Γ in the above equation denotes the gamma function. The parameters a, b, c, d correspond to the shape and scale parameters of two probability density functions of the gamma distribution (see spm_Gpdf.m), one corresponding to the main response and the other to the undershoot. The parameter c determines the ratio of the response to the undershoot. As described by the spm_hrf function in SPM12, the parameters a, b, c, and d are obtained by priors with different meanings (see more in spm_hrf.m). Taking into account the variability of the hemodynamic function across regions and voxels, these parameters are given latent variables, analogous to the approach of Zeidman et al ^25^, thus adapting them to the optimization algorithm in the tool. These parameters also describe the properties of white matter HRFs.

### 5.6. Bayesian Model Comparison

In the Bayesian pRF computational framework, each voxel is given a set of model evidence, which is used as a metric for model evaluation. We compared the evidence from the HCP data under six different pRF models using a random effects analysis (RFX) model with the free energy described in Zeidman et al. ^25,68^. Each model was given its Expected Probability and Protection Exceedance Probability (PXP). The former refers to the probability that random subject data will be produced by each model, while the latter represents the probability that each model will outperform all other models in a comparison.

### 5.7. FMA-Cortical Functional Connectivity and Gradients Cortical ROI Definition and Connectivity Matrix Computation

To parcellate the visual cortex, we defined a set of 36 functional regions of interest (ROIs). These ROIs were created by intersecting a probabilistic atlas of the visual cortex ^69^ with retinotopic maps generated by Neuropythy ^70^. The ROIs were organized on a grid spanning areas V1/V2 across four eccentricity bands ({0°, 2°, 8°, 20°}) and seven polar angle bins ({-30°, 30°, 90°, 150°, 210°, 270°, 330°}). Using the group-averaged retinotopy fMRI data, we extracted the mean time-series for each of the 36 cortical ROIs. A dense FMA-cortical functional connectivity matrix was then computed by calculating the Pearson correlation between each cortical ROI’s time-series and the individual time-series of every voxel in the FMA. The resulting correlation values were subsequently stabilized using a Fisher’s Z-transform.

### Gradient Computation and Cortical Projection

The principal axes of functional connectivity variation within the FMA were identified by applying diffusion map embedding to the FMA-cortical connectivity matrix, through the BrainSpace Toolbox ^71^. This method produces a set of gradients, with the first gradient (*G*) representing the primary, most dominant axis of connectivity change across all FMA voxels.

To visualize how the FMA’s primary functional gradient relates to the cortex, we projected it onto the cortical surface using a weighted summation approach adapted from prior work ^72^. The cortical projection map was calculated using the following formula:

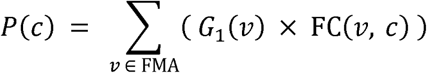

where *P*(*c*) is the projection value at a cortical vertex *c*, *G*(*v*) is the score of an FMA voxel v on the principal gradient, and *FC*(*v, c*) is the functional connectivity between them. Conceptually, this formula weights the cortical connectivity pattern of each FMA voxel by its score on the gradient. By summing these weighted maps, the resulting cortical projection *P*(*c*) reveals the two opposing patterns of cortical connectivity that define the FMA’s primary axis of functional organization.

### 5.8. Model Reliability Test

To evaluate the reliability of pRF estimates, we performed split-half analyses on group-averaged data across subjects. The 178 subjects were randomly divided into two groups of equal size, and the data were averaged within each group. Voxels were thresholded under the single pRF model, and polar angle, pRF size, and HRF parameters were estimated independently for each half.

### 5.9. Fiber Tracking

All tractography was performed in DSI Studio (http://dsistudio.labsolver.org/) utilizing the group-averaged Human Connectome Project (HCP-1065) template ^27^. A deterministic fiber tracking algorithm based on quantitative anisotropy was employed to generate streamlines ^26^. Tracking was seeded from all voxels within the FMA mask. To ensure comprehensive reconstruction of the fiber architecture, tracking was performed iteratively using a range of angular thresholds (30°, 40°, 50°, 60°, 70°, 80°, and 90°). Additional tracking parameters were held constant: a fractional anisotropy (FA) threshold of 0.15 was used to terminate pathways, the step size was set to 0.5 mm, and streamlines shorter than 20 mm or longer than 200 mm were discarded.

To delineate anatomical tracts corresponding to specific visual field representations, we employed a region-of-interest (ROI) and region-of-avoidance (ROA) strategy. The ROIs and ROAs were derived directly from our pRF functional maps, and the procedure was conducted separately for the left and right visual field pathways. For the left visual field pathway, a single seed volume (ROI) was created from all voxels functionally identified as representing the left visual field (in both hemispheres), while an avoidance volume (ROA) was created from all voxels representing the right visual field. Streamlines were discarded if they entered the avoidance region. An analogous seeding and avoidance strategy was applied to delineate the right visual field pathway. For each pathway (left and right visual field), the streamlines generated across all angular thresholds were merged into a single comprehensive bundle. To remove anatomically implausible or spurious fibers, these final bundles were refined by removing any streamline clusters containing fewer than 100 streamlines.

### 5.10. Neuronal Viral Tracing

#### Animal and viruses

Male C57BL/6J mice aged 3-5 months obtained from SPF (Beijing) Biotechnology Co., Ltd were used for this study. The recombinant adeno-associated viruses (rAAV-hSyn-EGFP (AAV/9, 5.01E+12vg/ml) and rAAV-hSyn-tdTomato (AAV/9, 3.75E+12vg/ml)) were purchased from BrainCase Co., Ltd. All animal experiments were approved by the Animal Care and Use Committee of the University of Science and Technology of China (USTCACUC21240122009).

#### Viral labeling

Viral injections were performed on anesthetized mice, which were subsequently mounted in an animal stereotaxic frame (RWD). A heating pad was utilized to maintain the animals’ core body temperature at 36 °C. For anterograde tracing, specific viral constructs were bilaterally delivered into the primary visual cortex (V1) of C57BL/6J male mice. The injection coordinates were established as: AP, −4.0 mm; ML, ±2.6 mm; and DV, −1.2 mm. Specifically, rAAV-hSyn-EGFP was infused into the right V1, and rAAV-hSyn-tdTomato into the left V1. Each injection consisted of 90 nL of virus, delivered via calibrated glass microelectrodes connected to an infusion pump at a constant rate of 10 nL/min. To minimize virus backflow along the injection track, the microsyringe was left in situ for 10 min following injection. Upon completion, the incision was sutured and the surgical site was sterilized with iodine.

#### Sample preparation and whole brain imaging

Three weeks post-viral injection, mice were anesthetized and transcardially perfused with 20 ml of 1×PBS (phosphate-buffered saline) twice, followed by 20 ml of 4% PFA (paraformaldehyde). Brains were then post-fixed in 4% PFA at 4 °C overnight and washed three times with 1×PBS. For hydrogel embedding, samples were immersed in a 1:1 solution of 4% HMS (Hydrogel Monomer Solution) and 20% BSA (bovine serum albumin). This solution was degassed in a vacuum pump for 10 min while centrifuge tubes were surrounded by ice. Samples were then polymerized in this solution at 37 °C for 4 hrs. All samples were subsequently sectioned into 300-μm-thick slices using a vibratome.

All slices underwent clearing in 4% SDS (Sodium Dodecyl Sulfate) with continuous shaking at 37 °C overnight. Following clearing, slices were washed three times with 0.3% PBST (1×PBS with 0.3% Triton X-100) and stored in 1×PBS at 4 °C. The processed slices were then sequentially mounted onto hydrophilic-treated glass panes. After applying vacuum-degassed 4% HMS solution in a fume hood, slices were covered with hydrophobic-treated glass slides. These samples were transferred to a 37 °C incubator and polymerized for 4 hrs. Subsequently, the samples were immersed in a high-refractive-index matching solution. The cleared slices were then imaged using VISoR (Volumetric Imaging with Synchronized on-the-fly-scan and Readout). Imaging was performed sequentially using 488 nm and 561 nm excitation for virus-labeled signal acquisition, and 405 nm excitation to capture tissue autofluorescence. Image stacks were acquired at an isotropic voxel size of 1×1×3.5 μm^3^.

#### Whole brain reconstruction

The brain reconstruction pipeline comprised five sequential stages ^28,34^: (1) Image columns are generated by the actual stacking of coordinates. Whole-brain slices were computationally reassembled from sequentially acquired image columns by applying pre-calibrated distortion correction coefficients and spatially registering overlapping fiducial markers to ground-truth coordinates. (2) The upper and lower image planes of each contrast-enhanced slice were algorithmically reconstructed through linear regression and interpolation. (3) Adjacent slices were co-registered using a hybrid approach that combines rigid transformation with B-spline-based non-rigid deformation to align fine-scale textural features and edge contours. (4) Displacement vector fields between neighboring slices were iteratively optimized to mitigate error propagation across the volumetric dataset. (5) A moving-least-squares algorithm was applied to minimize residual slice-level distortions while preserving intrinsic tissue morphology.

#### Quantification of spatial colocalization

To assess the degree of spatial colocalization between the two fluorescently labeled fiber populations, we calculated the Dice coefficient (D) and the Jaccard index (J) based on volumetric overlap of red and green channel-labeled projection. The metrics are defined as:

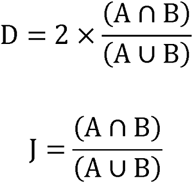

Where A and B denote the 3D volumes of green and red fiber tracts respectively, (A ∩ B) represents their volumetric overlap, and (A ∪ B) represents the combined volumetric space occupied by both tracts. The Jaccard index (J) and Dice coefficient (D) were calculated from binarized volumetric masks, with both metrics yielding values from 0 (complete spatial segregation) to 1 (perfect volumetric colocalization).

## 6. Data availability

The data and code will be shared upon requests.

## Supporting information

all supplemental figure

## Acknowledgements

This work was partially supported by National Science and Technology Innovation 2030 Major Program 2022ZD0204801, the Institute of Artificial Intelligence of Hefei Comprehensive National Science Center (23YGXT004, Y1022502).

## References

1. Gavrish, M., Kustova, A., Celis Suescún, J.C., Bessa, P., Mitina, N., and Tarabykin, V. (2024). Molecular mechanisms of corpus callosum development: a four-step journey. Frontiers in Neuroanatomy 17.

2. Suárez, R., Paolino, A., Fenlon, L.R., Morcom, L.R., Kozulin, P., Kurniawan, N.D., and Richards, L.J. (2018). A pan-mammalian map of interhemispheric brain connections predates the evolution of the corpus callosum. Proceedings of the National Academy of Sciences 115, 9622–9627. 10.1073/pnas.1808262115.

3. Gazzaniga, M.S. (2000). Cerebral specialization and interhemispheric communication: Does the corpus callosum enable the human condition? Brain 123, 1293–1326. 10.1093/brain/123.7.1293.

4. Schulte, T., and Müller-Oehring, E.M. (2010). Contribution of Callosal Connections to the Interhemispheric Integration of Visuomotor and Cognitive Processes. Neuropsychol Rev 20, 174–190. 10.1007/s11065-010-9130-1.

5. Wandell, B.A., Dumoulin, S.O., and Brewer, A.A. (2007). Visual field maps in human cortex. Neuron 56, 366–383. 10.1016/j.neuron.2007.10.012.

6. Bocci, T., Pietrasanta, M., Cerri, C., Restani, L., Caleo, M., and Sartucci, F. (2014). Visual callosal connections: role in visual processing in health and disease. Reviews in the Neurosciences 25, 113–127. 10.1515/revneuro-2013-0025.

7. Rivest, J., Cavanagh, P., and Lassonde, M. (1994). Interhemispheric depth judgement. Neuropsychologia 32, 69–76. 10.1016/0028-3932(94)90069-8.

8. Goldstein, A., Covington, B.P., Mahabadi, N., and Mesfin, F.B. (2025). Neuroanatomy, Corpus Callosum. In StatPearls (StatPearls Publishing).

9. Chung, S., Fieremans, E., Novikov, D.S., and Lui, Y.W. (2024). Microstructurally informed subject-specific parcellation of the corpus callosum using axonal water fraction. Brain Struct Funct 230, 1. 10.1007/s00429-024-02872-7.

10. Huang, H., Jiang, Y., Li, H., Wu, H., Feng, X., Gong, J., Jiang, S., Yao, D., and Luo, C. (2024). Functional organization of the human corpus callosum unveiled with BOLD-fMRI gradients. Imaging Neuroscience 2, 1–22. 10.1162/imag_a_00115.

11. Santana, C., Abreu, T., Rodrigues, J., Julio, P., Appenzeller, S., and Rittner, L. (2023). DTI-based Corpus Callosum parcellation using the Tensorial Morphological Gradient and Self-Organizing Maps. In 2023 19th International Symposium on Medical Information Processing and Analysis (SIPAIM), pp. 1–5. 10.1109/SIPAIM56729.2023.10373443.

12. Liu, P., Doehler, J., Henschke, J.U., Northall, A., Knaf-Serian, A., Loaiza-Carvajal, L.C., Budinger, E., Schwarzkopf, D.S., Speck, O., Pakan, J.M.P., et al. (2025). Layer-specific changes in sensory cortex across the lifespan in mice and humans. Nat Neurosci 28, 1978–1989. 10.1038/s41593-025-02013-1.

13. Yang, J., Huber, L., Yu, Y., and Bandettini, P.A. (2021). Linking cortical circuit models to human cognition with laminar fMRI. Neuroscience & Biobehavioral Reviews 128, 467–478. 10.1016/j.neubiorev.2021.07.005.

14. Yu, Y., Huber, L., Yang, J., Jangraw, D.C., Handwerker, D.A., Molfese, P.J., Chen, G., Ejima, Y., Wu, J., and Bandettini, P.A. (2019). Layer-specific activation of sensory input and predictive feedback in the human primary somatosensory cortex. Science Advances 5, eaav9053. 10.1126/sciadv.aav9053.

15. Rochefort, N.L., Buzás, P., Quenech’du, N., Koza, A., Eysel, U.T., Milleret, C., and Kisvárday, Z.F. (2009). Functional Selectivity of Interhemispheric Connections in Cat Visual Cortex. Cerebral Cortex 19, 2451–2465. 10.1093/cercor/bhp001.

16. Sincich, L.C., and Blasdel, G.G. (2001). Oriented Axon Projections in Primary Visual Cortex of the Monkey. J Neurosci 21, 4416–4426. 10.1523/JNEUROSCI.21-12-04416.2001.

17. Ding, Z., Huang, Y., Bailey, S.K., Gao, Y., Cutting, L.E., Rogers, B.P., Newton, A.T., and Gore, J.C. (2018). Detection of synchronous brain activity in white matter tracts at rest and under functional loading. Proceedings of the National Academy of Sciences 115, 595–600. 10.1073/pnas.1711567115.

18. Fraser, L.M., Stevens, M.T., Beyea, S.D., and D’Arcy, R.C.N. (2012). White versus gray matter: fMRI hemodynamic responses show similar characteristics, but differ in peak amplitude. BMC Neuroscience 13, 91. 10.1186/1471-2202-13-91.

19. Huang, Y., Wei, P.-H., Xu, L., Chen, D., Yang, Y., Song, W., Yi, Y., Jia, X., Wu, G., Fan, Q., et al. (2023). Intracranial electrophysiological and structural basis of BOLD functional connectivity in human brain white matter. Nat Commun 14, 3414. 10.1038/s41467-023-39067-3.

20. Ji, G.-J., Sun, J., Hua, Q., Zhang, L., Zhang, T., Bai, T., Wei, L., Wang, X., Qiu, B., Wang, A., et al. (2023). White matter dysfunction in psychiatric disorders is associated with neurotransmitter and genetic profiles. Nat. Mental Health 1, 655–666. 10.1038/s44220-023-00111-2.

21. Zhao, Y., Yang, Z., Ding, Z., and Su, J. (2023). A Riemannian Framework for Structurally Curated Functional Clustering of Brain White Matter Fibers. IEEE Transactions on Medical Imaging 42, 2414–2424. 10.1109/TMI.2023.3252269.

22. Wang, H., Wang, X., Wang, Y., Zhang, D., Yang, Y., Zhou, Y., Qiu, B., and Zhang, P. (2023). White matter BOLD signals at 7 Tesla reveal visual field maps in optic radiation and vertical occipital fasciculus. NeuroImage 269, 119916. 10.1016/j.neuroimage.2023.119916.

23. Wang, Y., Wang, H., Hu, S., Nguchu, B.A., Zhang, D., Chen, S., Ji, Y., Qiu, B., and Wang, X. (2024). Sub-bundle based analysis reveals the role of human optic radiation in visual working memory. Human Brain Mapping 45, e26800. 10.1002/hbm.26800.

24. Benson, N.C., Jamison, K.W., Arcaro, M.J., Vu, A.T., Glasser, M.F., Coalson, T.S., Van Essen, D.C., Yacoub, E., Ugurbil, K., Winawer, J., et al. (2018). The Human Connectome Project 7 Tesla retinotopy dataset: Description and population receptive field analysis. Journal of Vision 18, 23. 10.1167/18.13.23.

25. Zeidman, P., Silson, E.H., Schwarzkopf, D.S., Baker, C.I., and Penny, W. (2018). Bayesian population receptive field modelling. Neuroimage 180, 173–187. 10.1016/j.neuroimage.2017.09.008.

26. Yeh, F.C., Verstynen, T.D., Wang, Y., Fernandez-Miranda, J.C., and Tseng, W.Y. (2013). Deterministic diffusion fiber tracking improved by quantitative anisotropy. PLoS One 8, e80713. 10.1371/journal.pone.0080713.

27. Yeh, F.-C., Panesar, S., Fernandes, D., Meola, A., Yoshino, M., Fernandez-Miranda, J.C., Vettel, J.M., and Verstynen, T. (2018). Population-averaged atlas of the macroscale human structural connectome and its network topology. NeuroImage 178, 57–68. 10.1016/j.neuroimage.2018.05.027.

28. Ma, C., Xia, D., Huang, S., Du, Q., Liu, J., Zhang, B., Zhu, Q., Bi, G., Wang, H., and Xu, R.X. (2023). High precision vibration sectioning for 3D imaging of the whole central nervous system. Journal of Neuroscience Methods 399, 109966. 10.1016/j.jneumeth.2023.109966.

29. Wang, H., Zhu, Q., Ding, L., Shen, Y., Yang, C.-Y., Xu, F., Shu, C., Guo, Y., Xiong, Z., Shan, Q., et al. (2019). Scalable volumetric imaging for ultrahigh-speed brain mapping at synaptic resolution. National Science Review 6, 982–992. 10.1093/nsr/nwz053.

30. Li, M., Newton, A.T., Anderson, A.W., Ding, Z., and Gore, J.C. (2019). Characterization of the hemodynamic response function in white matter tracts for event-related fMRI. Nature Communications 10, 1140. 10.1038/s41467-019-09076-2.

31. Sporns, O., Tononi, G., and Kötter, R. (2005). The Human Connectome: A Structural Description of the Human Brain. PLOS Computational Biology 1, e42. 10.1371/journal.pcbi.0010042.

32. Van Essen, D.C., Smith, S.M., Barch, D.M., Behrens, T.E., Yacoub, E., Ugurbil, K., and W. U-Minn HCP Consortium (2013). The WU-Minn Human Connectome Project: an overview. NeuroImage 80, 62–79. 10.1016/j.neuroimage.2013.05.041.

33. Zingg, B., Hintiryan, H., Gou, L., Song, M.Y., Bay, M., Bienkowski, M.S., Foster, N.N., Yamashita, S., Bowman, I., Toga, A.W., et al. (2014). Neural networks of the mouse neocortex. Cell 156, 1096–1111. 10.1016/j.cell.2014.02.023.

34. Zheng, W., Mu, H., Chen, Z., Liu, J., Xia, D., Cheng, Y., Jing, Q., Lau, P.-M., Tang, J., Bi, G.-Q., et al. (2024). NEATmap: a high-efficiency deep learning approach for whole mouse brain neuronal activity trace mapping. National Science Review 11, nwae109. 10.1093/nsr/nwae109.

35. Oh, S.W., Harris, J.A., Ng, L., Winslow, B., Cain, N., Mihalas, S., Wang, Q., Lau, C., Kuan, L., Henry, A.M., et al. (2014). A mesoscale connectome of the mouse brain. Nature 508, 207–214. 10.1038/nature13186.

36. Azadbakht, H., Parkes, L.M., Haroon, H.A., Augath, M., Logothetis, N.K., de Crespigny, A., D’Arceuil, H.E., and Parker, G.J.M. (2015). Validation of High-Resolution Tractography Against In Vivo Tracing in the Macaque Visual Cortex. Cereb Cortex 25, 4299–4309. 10.1093/cercor/bhu326.

37. Schilling, K.G., Daducci, A., Maier-Hein, K., Poupon, C., Houde, J.C., Nath, V., Anderson, A.W., Landman, B.A., and Descoteaux, M. (2019). Challenges in diffusion MRI tractography - Lessons learned from international benchmark competitions. Magn Reson Imaging 57, 194–209. 10.1016/j.mri.2018.11.014.

38. Thomas, C., Ye, F.Q., Irfanoglu, M.O., Modi, P., Saleem, K.S., Leopold, D.A., and Pierpaoli, C. (2014). Anatomical accuracy of brain connections derived from diffusion MRI tractography is inherently limited. Proceedings of the National Academy of Sciences 111, 16574–16579. 10.1073/pnas.1405672111.

39. Ding, Z., Newton, A.T., Xu, R., Anderson, A.W., Morgan, V.L., and Gore, J.C. (2013). Spatio-Temporal Correlation Tensors Reveal Functional Structure in Human Brain. PLOS ONE 8, e82107. 10.1371/journal.pone.0082107.

40. Nozais, V., Forkel, S.J., Foulon, C., Petit, L., and Thiebaut de Schotten, M. (2021). Functionnectome as a framework to analyse the contribution of brain circuits to fMRI. Commun Biol 4, 1035. 10.1038/s42003-021-02530-2.

41. Suárez, R., Gobius, I., and Richards, L.J. (2014). Evolution and development of interhemispheric connections in the vertebrate forebrain. Front. Hum. Neurosci. 8. 10.3389/fnhum.2014.00497.

42. Fenlon, L.R., and Richards, L.J. (2015). Contralateral targeting of the corpus callosum in normal and pathological brain function. Trends in Neurosciences 38, 264–272. 10.1016/j.tins.2015.02.007.

43. Van Essen, D.C., Donahue, C.J., Coalson, T.S., Kennedy, H., Hayashi, T., and Glasser, M.F. (2019). Cerebral cortical folding, parcellation, and connectivity in humans, nonhuman primates, and mice. Proceedings of the National Academy of Sciences 116, 26173–26180. 10.1073/pnas.1902299116.

44. Parsons, N., Irimia, A., Amgalan, A., Ugon, J., Morgan, K., Shelyag, S., Hocking, A., Poudel, G., and Caeyenberghs, K. (2023). Structural-functional connectivity bandwidth predicts processing speed in mild traumatic brain Injury: A multiplex network analysis. NeuroImage: Clinical 38, 103428. 10.1016/j.nicl.2023.103428.

45. Lim, J.-S., and Kang, D.-W. (2015). Stroke Connectome and Its Implications for Cognitive and Behavioral Sequela of Stroke. J Stroke 17, 256–267. 10.5853/jos.2015.17.3.256.

46. Ji, G.-J., Cui, Z., D’Arcy, R.C.N., Liao, W., Biswal, B.B., Zhang, Q., Luo, C., Zang, Y.-F., Ding, Z., Zuo, X.-N., et al. (2025). Imaging brain white matter function using resting-state functional MRI. Science Bulletin 70, 1384–1388. 10.1016/j.scib.2024.11.001.

47. Wang, P., Wang, J., Michael, A., Wang, Z., Klugah-Brown, B., Meng, C., and Biswal, B.B. (2022). White Matter Functional Connectivity in Resting-State fMRI: Robustness, Reliability, and Relationships to Gray Matter. Cerebral Cortex 32, 1547–1559. 10.1093/cercor/bhab181.

48. Zu, Z., Choi, S., Zhao, Y., Gao, Y., Li, M., Schilling, K.G., Ding, Z., and Gore, J.C. (2024). The missing third dimension—Functional correlations of BOLD signals incorporating white matter. Science Advances 10, eadi0616. 10.1126/sciadv.adi0616.

49. Chu, T., Si, X., Song, X., Che, K., Dong, F., Guo, Y., Chen, D., Yao, W., Zhao, F., Xie, H., et al. (2025). Understanding structural-functional connectivity coupling in patients with major depressive disorder: A white matter perspective. Journal of Affective Disorders 373, 219–226. 10.1016/j.jad.2024.12.082.

50. Li, J., Biswal, B.B., Meng, Y., Yang, S., Duan, X., Cui, Q., Chen, H., and Liao, W. (2020). A neuromarker of individual general fluid intelligence from the white-matter functional connectome. Transl Psychiatry 10, 147. 10.1038/s41398-020-0829-3.

51. Huber, L. (Renzo), Kronbichler, L., Stirnberg, R., Ehses, P., Stöcker, T., Fernández-Cabello, S., Poser, B.A., and Kronbichler, M. (2023). Evaluating the capabilities and challenges of layer-fMRI VASO at 3T. Apert Neuro 3, 10.52294/001c.85117. http://doi.org/10.52294/001c.85117.

52. Breau, M.A., and Trembleau, A. (2023). Chemical and mechanical control of axon fasciculation and defasciculation. Semin Cell Dev Biol 140, 72–81. 10.1016/j.semcdb.2022.06.014.

53. Moreland, T., and Poulain, F.E. (2022). To Stick or Not to Stick: The Multiple Roles of Cell Adhesion Molecules in Neural Circuit Assembly. Front. Neurosci. 16. 10.3389/fnins.2022.889155.

54. Yan, K., Bormuth, I., Bormuth, O., Tutukova, S., Renner, A., Bessa, P., Schaub, T., Rosário, M., and Tarabykin, V. (2023). TrkB-dependent EphrinA reverse signaling regulates callosal axon fasciculate growth downstream of Neurod2/6. Cereb Cortex 33, 1752–1767. 10.1093/cercor/bhac170.

55. Tezuka, Y., Hagihara, K.M., Ohki, K., Hirano, T., and Tagawa, Y. (2022). Developmental stage-specific spontaneous activity contributes to callosal axon projections. eLife 11, e72435. 10.7554/eLife.72435.

56. Wandell, B.A., and Winawer, J. (2015). Computational neuroimaging and population receptive fields. Trends in cognitive sciences 19, 349–357. 10.1016/j.tics.2015.03.009.

57. Caleo, M., Innocenti, G.M., and Ptito, M. (2013). Physiology and Plasticity of Interhemispheric Connections. Neural Plast 2013, 176183. 10.1155/2013/176183.

58. Foubert, L. (2007). Spatio-temporal characteristics of the visual interhemispheric integration via the corpus callosum: computational modeling & optical imaging approaches.

59. Schmidt, K.E., Lomber, S.G., and Innocenti, G.M. (2010). Specificity of Neuronal Responses in Primary Visual Cortex Is Modulated by Interhemispheric CorticoCortical Input. Cereb Cortex 20, 2776–2786. 10.1093/cercor/bhq024.

60. Glasser, M.F., Sotiropoulos, S.N., Wilson, J.A., Coalson, T.S., Fischl, B., Andersson, J.L., Xu, J., Jbabdi, S., Webster, M., Polimeni, J.R., et al. (2013). The minimal preprocessing pipelines for the Human Connectome Project. Neuroimage 80, 105–124. 10.1016/j.neuroimage.2013.04.127.

61. T. Vu, A., Jamison, K., Glasser, M.F., Smith, S.M., Coalson, T., Moeller, S., Auerbach, E.J., Uğurbil, K., and Yacoub, E. (2017). Tradeoffs in pushing the spatial resolution of fMRI for the 7T Human Connectome Project. NeuroImage 154, 23–32. 10.1016/j.neuroimage.2016.11.049.

62. Avants, B.B., Epstein, C.L., Grossman, M., and Gee, J.C. (2008). Symmetric diffeomorphic image registration with cross-correlation: Evaluating automated labeling of elderly and neurodegenerative brain. Medical Image Analysis 12, 26–41. 10.1016/j.media.2007.06.004.

63. Jenkinson, M., Beckmann, C.F., Behrens, T.E.J., Woolrich, M.W., and Smith, S.M. (2012). FSL. NeuroImage 62, 782–790. 10.1016/j.neuroimage.2011.09.015.

64. Radwan, A.M., Sunaert, S., Schilling, K., Descoteaux, M., Landman, B.A., Vandenbulcke, M., Theys, T., Dupont, P., and Emsell, L. (2022). An atlas of white matter anatomy, its variability, and reproducibility based on constrained spherical deconvolution of diffusion MRI. NeuroImage 254, 119029. 10.1016/j.neuroimage.2022.119029.

65. Smith, S.M., Jenkinson, M., Woolrich, M.W., Beckmann, C.F., Behrens, T.E.J., Johansen-Berg, H., Bannister, P.R., De Luca, M., Drobnjak, I., Flitney, D.E., et al. (2004). Advances in functional and structural MR image analysis and implementation as FSL. NeuroImage 23, S208–S219. 10.1016/j.neuroimage.2004.07.051.

66. Boynton, G.M., Engel, S.A., Glover, G.H., and Heeger, D.J. (1996). Linear systems analysis of functional magnetic resonance imaging in human V1. J Neurosci 16, 4207–4221. 10.1523/JNEUROSCI.16-13-04207.1996.

67. Friston, K.J., Fletcher, P., Josephs, O., Holmes, A., Rugg, M.D., and Turner, R. (1998). Event-related fMRI: characterizing differential responses. Neuroimage 7, 30–40. 10.1006/nimg.1997.0306.

68. Stephan, K.E., Penny, W.D., Daunizeau, J., Moran, R.J., and Friston, K.J. (2009). Bayesian model selection for group studies. Neuroimage 46, 1004–1017. 10.1016/j.neuroimage.2009.03.025.

69. Wang, L., Mruczek, R.E., Arcaro, M.J., and Kastner, S. (2015). Probabilistic Maps of Visual Topography in Human Cortex. Cerebral Cortex 25, 3911–3931. 10.1093/cercor/bhu277.

70. Benson, N.C., and Winawer, J. (2018). Bayesian analysis of retinotopic maps. eLife 7. 10.7554/eLife.40224.

71. Vos de Wael, R., Benkarim, O., Paquola, C., Lariviere, S., Royer, J., Tavakol, S., Xu, T., Hong, S.-J., Langs, G., Valk, S., et al. (2020). BrainSpace: a toolbox for the analysis of macroscale gradients in neuroimaging and connectomics datasets. Commun Biol 3, 1–10. 10.1038/s42003-020-0794-7.

72. Katsumi, Y., Zhang, J., Chen, D., Kamona, N., Bunce, J.G., Hutchinson, J.B., Yarossi, M., Tunik, E., Dickerson, B.C., Quigley, K.S., et al. (2023). Correspondence of functional connectivity gradients across human isocortex, cerebellum, and hippocampus. Commun Biol 6, 401. 10.1038/s42003-023-04796-0.

