## Supplementary material for "Parallel Wires: A Conserved Principle of Contralateral-Ipsilateral Segregation in the Visual Corpus Callosum": all supplemental figure

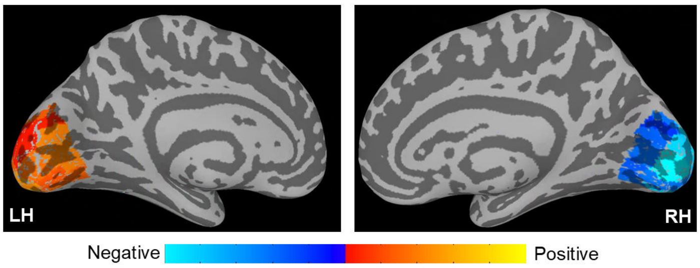


**Fig. S1. Cortical Projection of the FMA Functional Gradient.** The principal functional gradient of the forceps major (FMA) is projected onto the visual cortex. The map validates the gradient by recovering the canonical contralateral organization of vision: positive values on the left hemisphere (LH) represent its strong functional link to the right visual field, while negative values on the right hemisphere (RH) represent its link to the left visual field in the Fig. 2B.

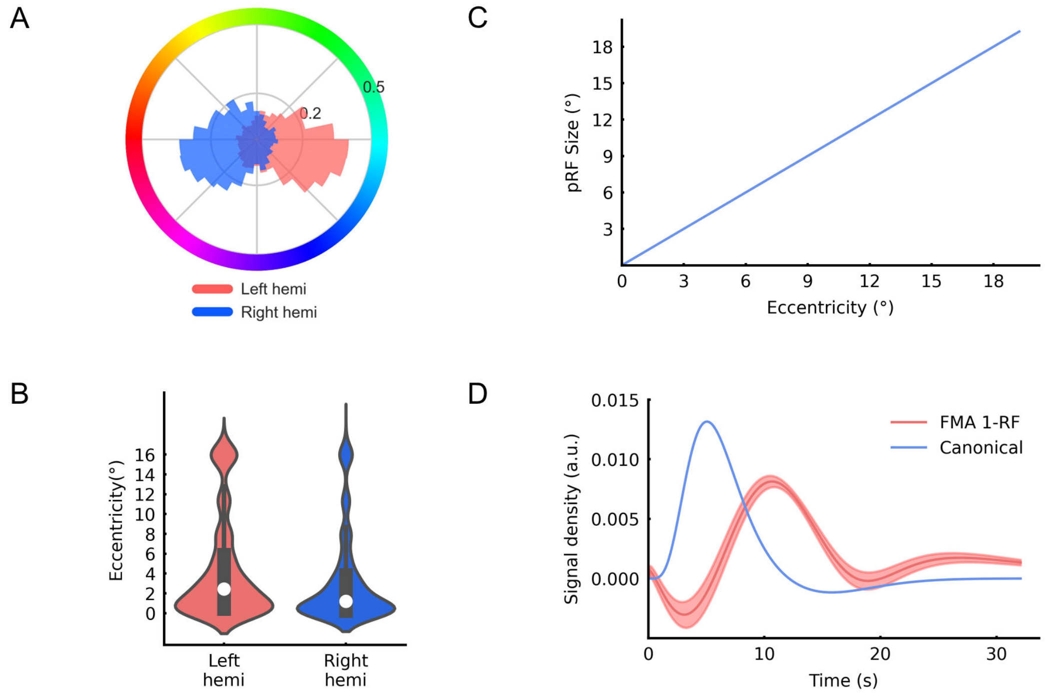


**Fig. S2. Visual field map properties, hemodynamic response functions (HRFs), and gradient-based validation of angular characteristics in the FMA fiber bundle. (A)** Polar angle distribution maps for the FMA, with voxels from the right hemisphere shown in blue and those from the left hemisphere in pink. **(B)** Violin plots showing the distribution of pRF eccentricity (in degrees of visual angle) within the FMA; white dots indicate the median values. **(C)** Relationship between pRF size (σ, in degrees of visual angle) and eccentricity. Solid lines indicate fitted linear regression models, and shaded areas denote bootstrapped 95% confidence intervals. **(D)** Fitted HRF curves for the FMA. Solid lines represent the mean HRF across voxels, and shaded areas represent bootstrapped 95% confidence intervals.

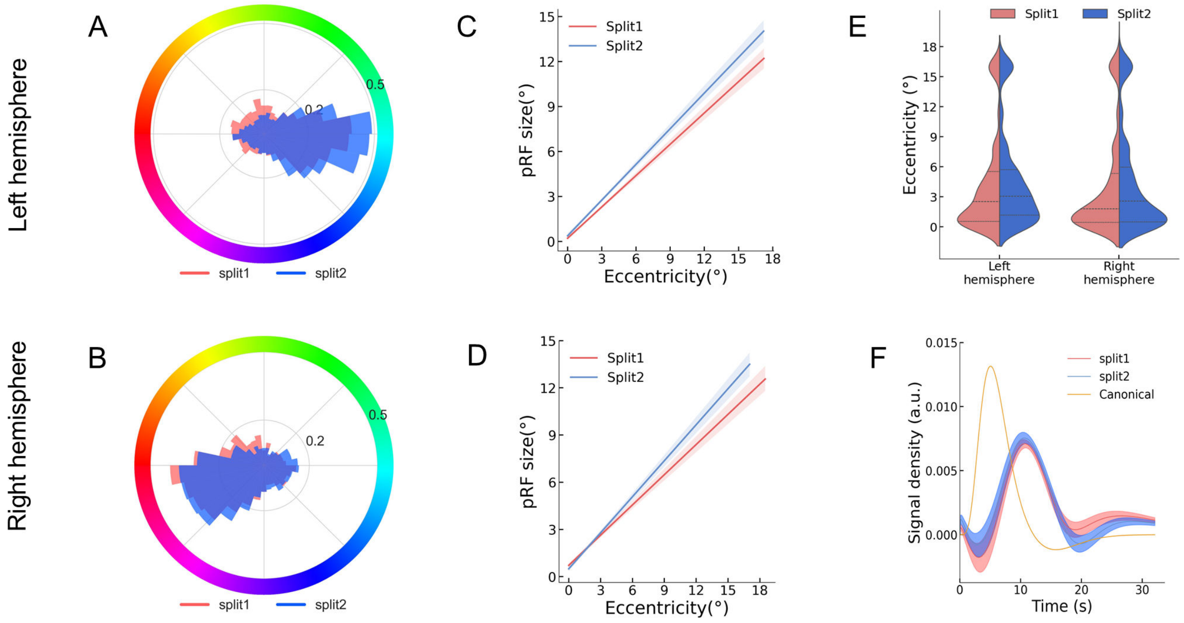


**Fig. S3. Results of split-half analyses across subjects using group-averaged data. (A, B)** Comparison of the polar angle distribution of FMA estimated from the data of two halves of subjects. Split 1 and 2 are represented in pink and blue, respectively. **(C, D)** The correlation between pRF size and eccentricity in the left and right hemispheres. **(E)** Eccentricity distribution and **(F)** HRFs in FMA for the split-subjects analysis. Shaded areas represent bootstrapped 95% confidence intervals.

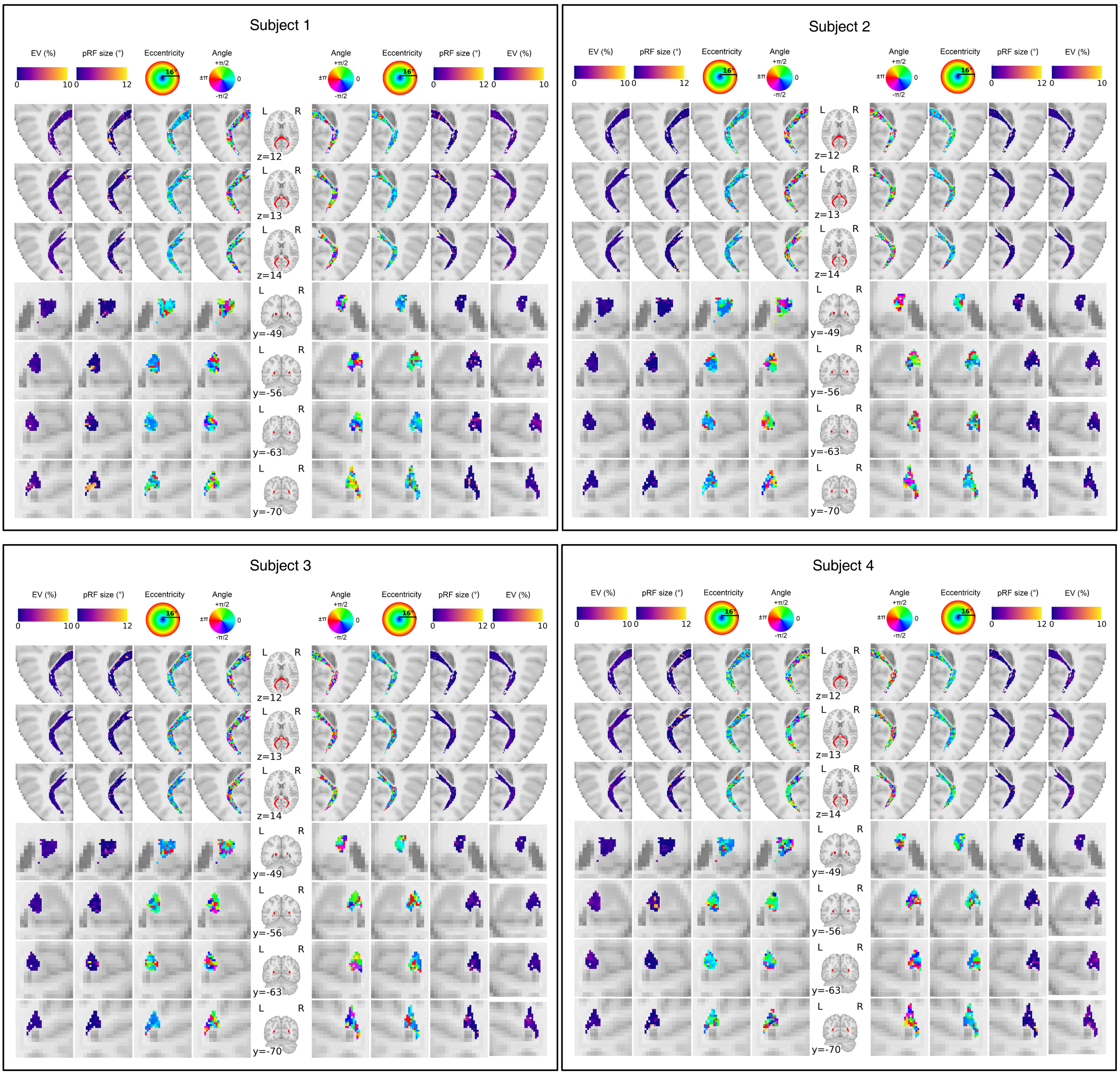


**Fig. S4. Individual-level pRF estimates from four randomly selected subjects.** Visual field maps and explained variance maps are shown in selected axial and coronal views, with all voxels within the region of interest (ROI) displayed. Columns, from the center to the outer sides, display: (i) the FMA fiber bundle mask, (ii) polar angle maps (in degrees of visual angle), (iii) eccentricity maps (in degrees of visual angle), (iv) pRF size (σ, in degrees of visual angle), and (v) explained variance (%). Rows 2–4 (from top to bottom) correspond to axial slices from dorsal to ventral; rows 5–7 (from top to bottom) correspond to coronal slices from anterior to posterior. All maps are overlaid on the ICBM 152 2009c standard brain template, with Montreal Neurological Institute (MNI) coordinates for the z-axis (axial views) and y-axis (coronal views) shown in millimeters.

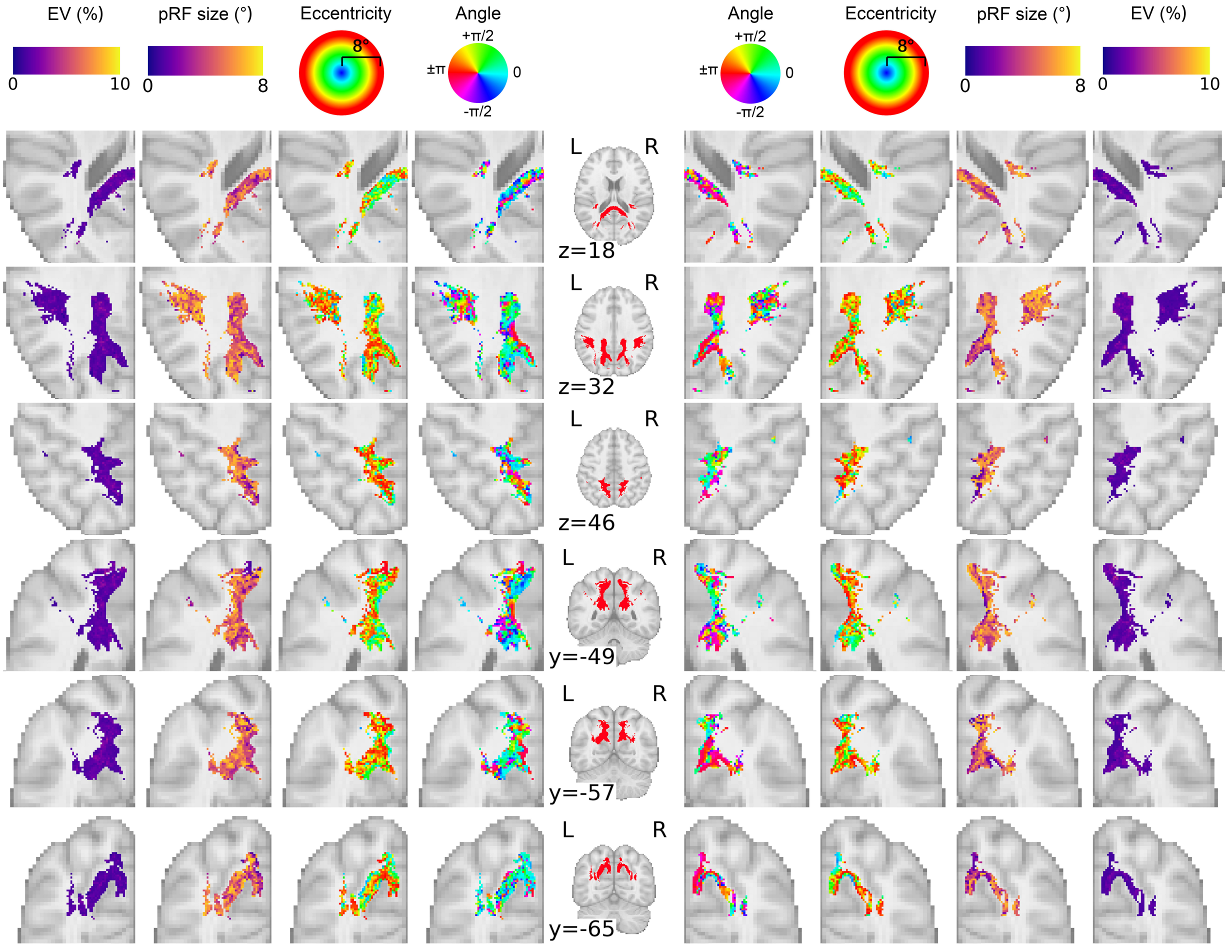


**Fig. S5. Population receptive field (pRF) parameter maps in the parietal corpus callosum (CC) connection fiber bundle.** Visual field maps and explained variance maps are shown in selected axial and coronal views, with all voxels within the region of interest (ROI) displayed. Columns, from the center to the outer sides, display: (i) the parietal CC connection fiber bundle mask, (ii) polar angle maps (in degrees of visual angle), (iii) eccentricity maps (in degrees of visual angle), (iv) pRF size (σ, in degrees of visual angle), and (v) explained variance (%). Rows 2–4 (from top to bottom) correspond to axial slices from dorsal to ventral; rows 5–7 (from top to bottom) correspond to coronal slices from anterior to posterior. All maps are overlaid on the ICBM 152 2009c standard brain template, with Montreal Neurological Institute (MNI) coordinates for the z-axis (axial views) and y-axis (coronal views) shown in millimeters.

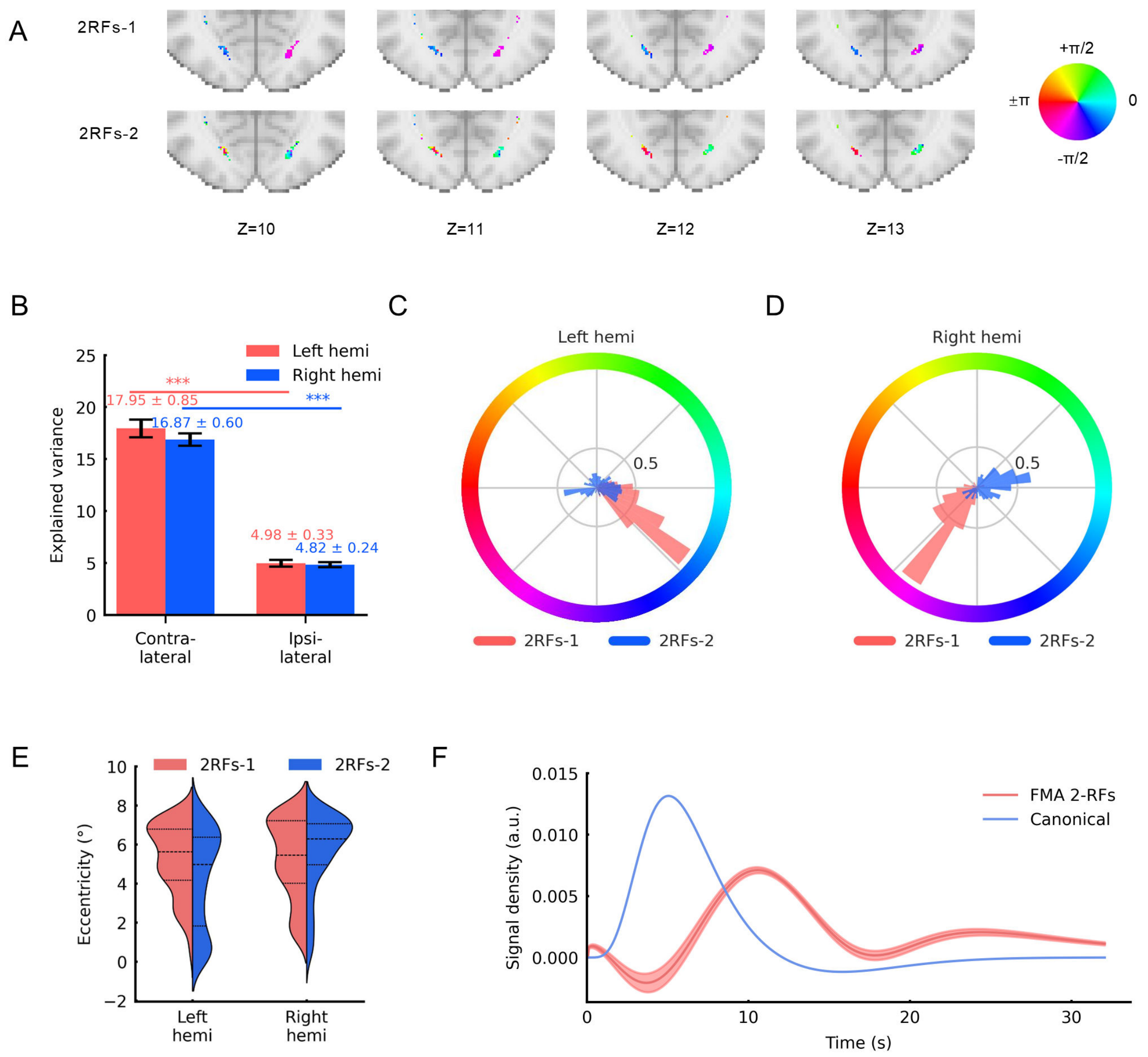


**Fig. S6. Polar angle distribution, visual field map properties and HRFs in dual pRF model winning voxels. (A)** Polar angle distribution maps from pRF estimation for dual receptive fields. The first row corresponds to pRF-1, and the second row to pRF-2 in the model. Slice position is indicated by the *z*-coordinate in MNI space (in millimeters). **(B)** Comparison of explained variance (%) between contralateral and ipsilateral voxels in the left and right hemispheres. A strong contralateral bias was evident: contralateral pRFs explained significantly more variance than their ipsilateral counterparts in both the left (17.95±0.85% vs. 4.98±0.33%, $p < 0.001$) and right hemispheres (16.87±0.60% vs. 4.82±0.24%, $p < 0.001$) **(C, D)** Normalized polar angle rectangular distribution maps for dual receptive fields in the left and right hemispheres, respectively; shaded area indicates overlapping angular ranges between pRF-1 and pRF-2. **(E)** Violin plots showing eccentricity distributions (in degrees of visual angle) in the left and right hemispheres for dual pRFs. Black dashed lines indicate the quartile boundaries in the violin plots. **(F)** Fitted HRF curves. Solid lines represent the mean HRF curve across voxels, and shaded areas represent bootstrapped 95% confidence intervals.

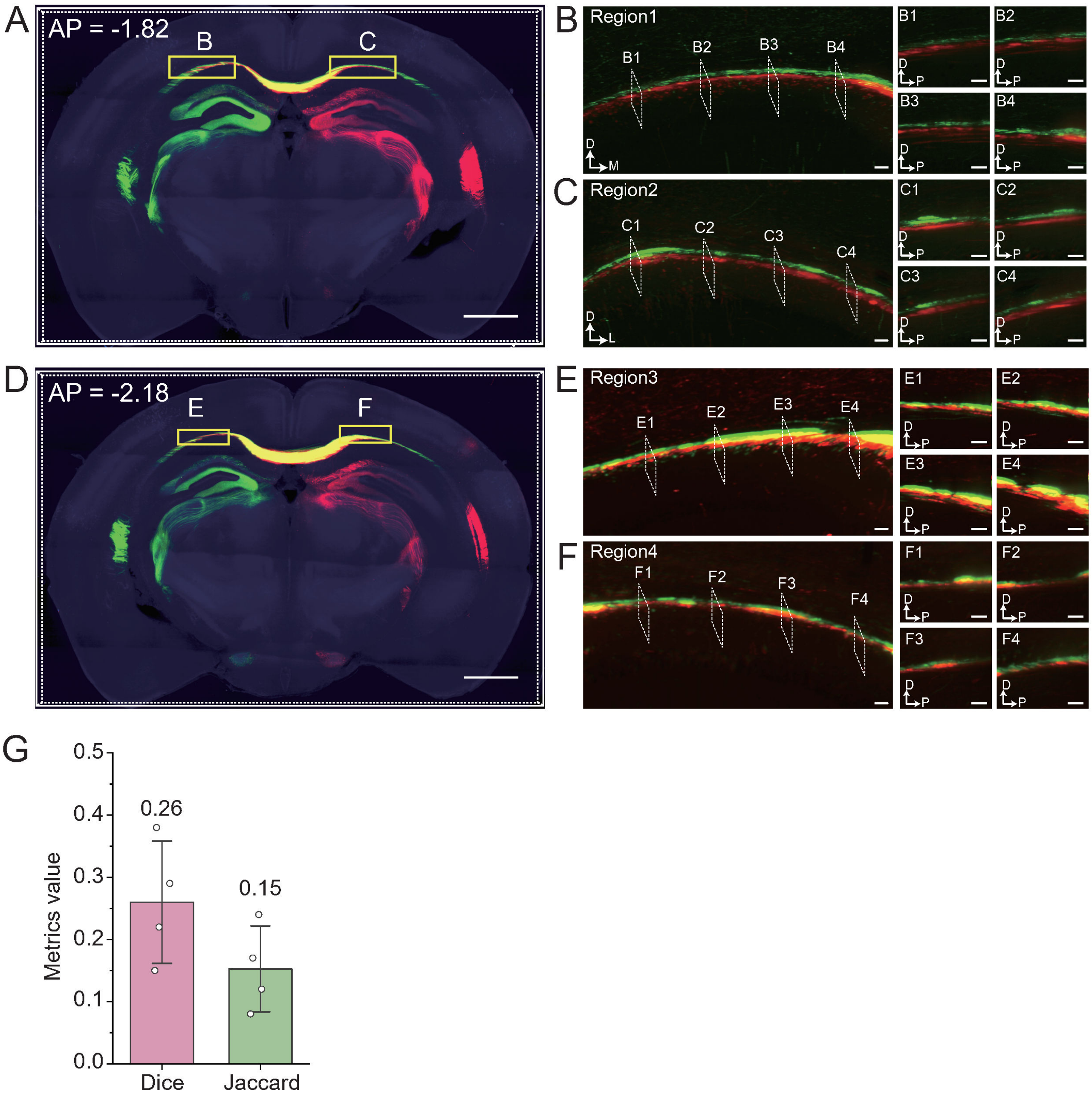


**Fig. S7**. **Quantification of spatial segregation in the mouse visual corpus callosum. (A, D)** Coronal views at two anterior-posterior (AP) levels. Note the transient zone of axonal overlap (yellow) at the commissural midline, which gives way to clear spatial segregation of the red and green fiber bundles more laterally. This provides a direct anatomical correlate for the sparse, localized dual-pRF voxels (midline integration) and the dominant single-pRF voxels (lateral segregation). Scale bar: 500 μm. **(B, C, E, F)** High-magnification analysis of the regions outlined in (A) and (D). The main panels show coronal views, while the corresponding sagittal sections (e.g., B1-B4) reveal a consistent laminar segregation. Axons from one hemisphere are consistently organized dorsal to axons from the other. This precise layered arrangement provides the structural mechanism that maintains the separation of the parallel pathways after they cross the midline. Scale bars: 50 μm. **(G)** Quantification of spatial overlap using the mean Dice and Jaccard indices for the four lateral regions. Low mean coefficient values (< 0.4) provide quantitative confirmation of the high degree of spatial segregation between the two hemispheric pathways outside of the immediate midline, consistent with a predominantly segregated functional architecture.
